# A comprehensive pipeline for accurate annotation and quantification of circRNAs

**DOI:** 10.1101/2019.12.15.876755

**Authors:** Avigayel Rabin, Reut Ashwal-Fluss, Shlomo Shenzis, Daniella Apelblat, Sebastian Kadener

**Affiliations:** Biological Chemistry Department, Silberman Institute of Life Sciences, The Hebrew University of Jerusalem, Jerusalem, 91904, Israel; Biology Department, Brandeis University, Waltham, MA, 02454, USA

**Keywords:** circular RNA, circRNAs, RNA metabolism, splicing, pipeline

## Abstract

Identification and quantification of circular RNAs (circRNAs) depends strongly on the utilized computational pipeline. Here we describe an integrative approach for accurate annotation and quantification of circRNAs. First, we utilize several circRNA-identification pipelines to annotate circRNAs in a given organism. Second, we build a short sequence index that is used to search the unaligned RNA-seq reads. Our approach allows full annotation of circRNAs with fewer false positives and negatives than any individual pipeline or combination of them. Moreover, our approach is more sensitive than any individual pipeline and allows more accurate quantification and larger number of differentially expressed circRNAs.

## BACKGROUND

Circular RNAs (circRNAs) are abundant RNAs generated through circularization of specific exons by a process called backsplicing [1–4]. As covalently closed circles, circRNAs are generally more stable than linear RNA transcripts. This is likely due to the lack of free ends that can be targeted by exonucleases. circRNAs have been found in bacteria, archaea, and most eukaryotes [5]. They are highly expressed in metazoans, particularly in the central nervous system (CNS) [1, 6–8]. Interestingly, circRNAs accumulate in the CNS as animals age in flies, worms, and mice [9, 10]. When first discovered, circRNAs were thought to be a byproduct of splicing; however, multiple studies in the past five years have clearly shown that at least some of these RNAs are functional. Two pioneering works showed that circRNAs can bind to and likely modulate miRNA function [11, 12]. Recent studies showed that these molecules can also regulate the activity of RNA binding proteins [13] and ribosome biogenesis [14] and that a subset of them encode proteins [15–17]. Their functionality has also been demonstrated *in vivo* [15, 18, 19], and there is evidence that circRNAs are disease biomarkers [20].

Many computational pipelines exist for *de novo* discovery and quantification of circRNAs from RNA-seq data [21–23]. These include Acfs [24], DCC [25], segemehl [26], CIRCexplorer [27], KNIFE [28], MapSplice2 [29], circRNA_finder [30], CIRI [31], and find_circ [11]. The pipelines differ in sensitivity, precision, runtime, and storage requirements [21] as shown in several independent studies [21, 32, 33]. Analysis of a large number of datasets allowed researchers to generate different circRNA databases [34]. One of the most popular circRNAs databases is known as circBASE [35], which contains annotations and information on many, but not all, circRNAs. Since many circRNAs have been already discovered, the very time-consuming [21] process of *de novo* identification of circRNAs for every RNA-seq library is redundant. Moreover, most circRNAs are identified by only subsets or only one pipeline, making it difficult to determine whether these are real circRNAs or sequencing or annotation artifacts. Therefore, relying on the results of only one pipeline for circRNA annotation and quantification is highly problematic [33].

Generally, circRNA detection relies on the identification of a splicing junction that is unique for a given circRNA (i.e., the backsplice junction). Importantly, other biological processes and technical artifacts can generate splicing junctions that can be confused with those characteristic of circRNAs [23]. These include splicing errors, trans-splicing, linear concatamers (i.e., from exon duplications), and artifacts resulting from the template switching activity of the reverse transcriptase [23]. Hence, circRNAs need to be validated experimentally. To do so, researchers use the 3’ exonuclease RNaseR, which efficiently degrades linear RNA sequences and does not affect circular RNA molecules [36]. Usually, the abundance of specific splicing junctions (determined by RT-PCR or RNA-seq) in samples pretreated with RNaseR is compared to that in samples not treated with the exonuclease. This strategy is useful for validation of circRNAs, but it is not reliable for quantification.

Here, we developed an alternative approach for both identifying and quantifying circRNAs. We first generate a reliable database of circRNAs by analyzing RNA-seq data from RNaseR and mock-treated samples through several existing pipelines. In the second step, we use this database to create a reference to which the RNA-seq data is aligned. We showed that our approach outperforms the use of single pipelines with regards to both annotation and quantification.

## RESULTS

### General approach

To accurately annotate and quantify circRNAs, we utilize a two-step approach that we call the Short Read CircRNA Pipeline (SRCP). The first step consists of the annotation of validated circRNAs. To do this, we utilize several pipelines to identify all putative circRNAs in two different RNA-seq libraries: one generated from total rRNA-depleted RNA and one generated from the same RNA pretreated with RNaseR. Comparison allows us to generate a database containing all validated circRNAs based on RNaseR resistance. Creating this database is time consuming, but it is only performed once for each type of sample (e.g., tissue and/or organism). Following the annotation of circRNAs, our pipeline generates a reference using bowtie2 of the back-splicing junctions and the canonical (linear) splicing junctions of the first and last exon contained in each circRNAs; the latter is used to quantify the linear RNA generated from the circRNA-hosting genes. We then align the RNA-seq reads to this reference using bowtie2 to identify circRNAs reads in the datasets. Finally, we align the back-splice junction reads to the genome and transcriptome and eliminate those that align to the genome or to the transcriptome. This is done in order to remove reads that cannot be unequivocally assigned to the circRNAs (Figure 1).

**Figure 1.**
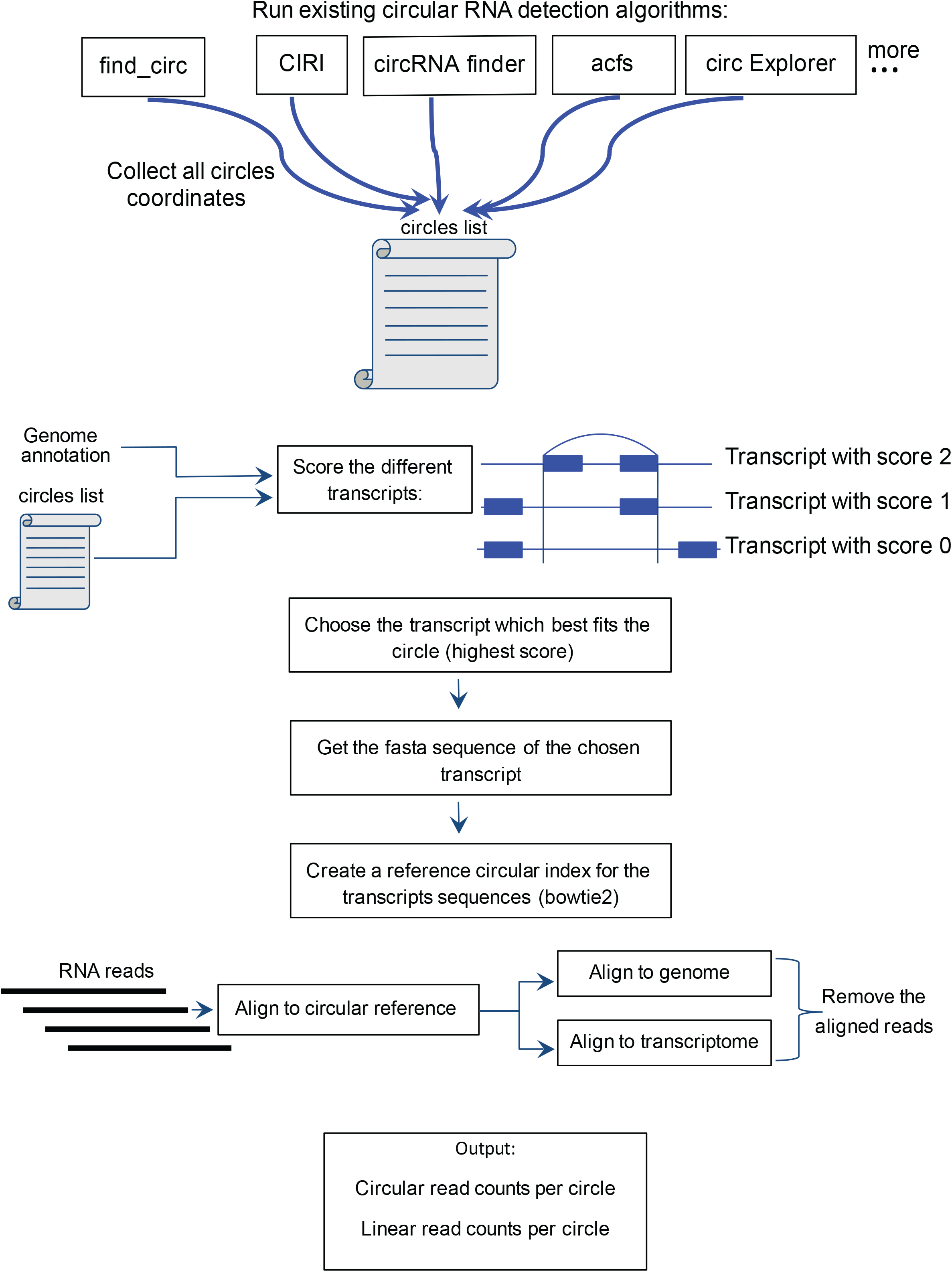
A comprehensive approach for annotation and quantification of circRNAs. The workflow of SRCP involves indexing of circRNA junctions identified using multiple circRNA detection pipelines. The reads are aligned to the genome and transcriptome and then to the circRNA index.

### No current pipeline accurately annotates all circRNAs

We began by determining what percentages of circRNAs in a previously published *Drosophila melanogaster* RNA-seq dataset obtained on samples with and without RNaseR treatment [GSE55872] were detected by five commonly used circRNA pipelines (find_circ [11], CIRI[31], CIRI2 [37], Acfs [24], circExplorer [27], and circRNA_finder [30]). As previously reported [21, 33], only about 10% of circRNAs were identified by all pipelines (Figure 2A). It has been previously proposed that a large fraction of pipeline-specific circRNAs are false positives and that circRNAs detected by several pipelines tend to be *bona fide* circRNAs [5, 33]. More than 50% of the detected circRNAs are detected by only one pipeline (Figure 2A).

**Figure 2.**
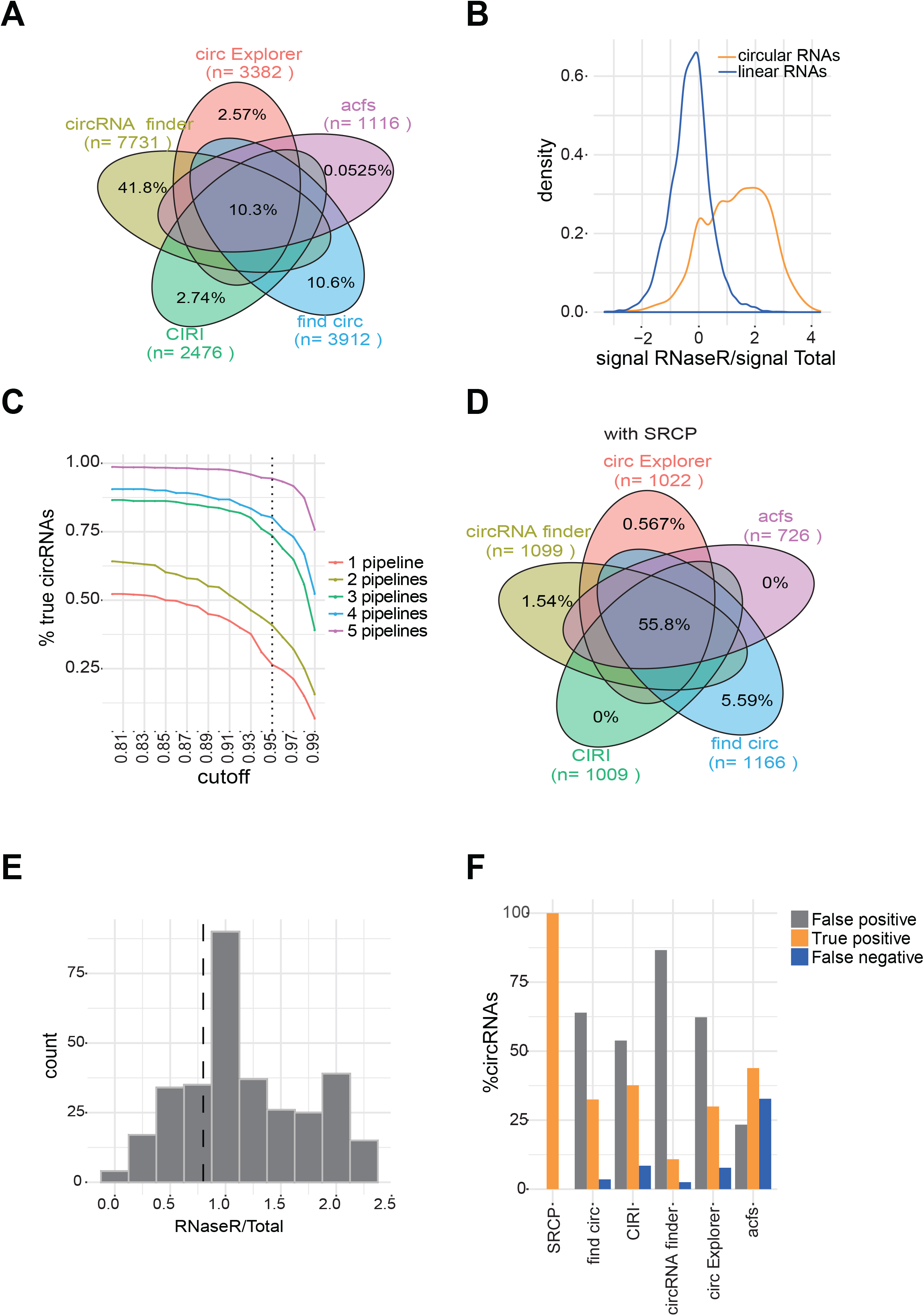
SRCP accurately annotates circRNAs. **A.**Venn diagram of the circRNAs found by the circRNA-identification pipelines in analysis of the total RNA library from the GSE55872 dataset. **B.** RNaseR/Total ratio distribution in *Drosophila melanogaster*. The data in orange represent the circular junctions and that in blue the linear junctions. **C.** The percent of circRNAs identified as “true” positives as a function of the cut-off for circRNAs identified by 1, 2, 3, 4, or 5 of the pipelines used. **D.** Venn diagram of the circRNAs found by the circRNA-identification pipelines using the SRCP approach. **E.** RNaseR/Total ratio of the pipeline-specific false-positive circRNAs. The dashed line indicates the cutoff value. All circles in the right of the line are annotated as “true” and the circles in the left as “false”. **F.** Percent false positives, false negatives, and true positives identified by SCRP and each individual circRNA-identification pipeline.

### Determination of false positives using RNAseR-seq datasets

The analyzed dataset contains RNA-seq reads from RNaseR-treated samples, which allowed us to determine with certainty that the identified circRNAs were real and not computational and/or methodological artifacts. For each identified circRNA junction we determined the ratio between the signal obtained in the RNaseR-treated samples and the signal in the total RNA (RNaseR/Total ratio). For each circRNA candidate, the larger the RNaseR/Total ratio, the more likely it represents a *bona fide* circRNA.

As expected, linear RNA junctions had lower RNAseR/Total RNA ratios than circRNAs junctions (Figure 2B). For linear mRNAs, the RNaseR/Total ratios have a discrete peak with a normal distribution. Interestingly, the putative circRNAs distribution is broader. This is likely due to overlapping peaks observed in the distribution of the candidate circRNAs. Therefore, it was necessary to apply a cut-off in order to distinguish *bona fide* circRNAs from false positives. Since false-positive circRNAs are linear molecules, we assumed that the distribution of the linear RNAs and the false-positive circRNAs should be similar. Therefore, we defined the error level to be an arbitrary percentage of the area on the right tail of the linear distribution. We accepted all circRNAs that have a higher RNaseR/Total ratio than this cut-off ratio as real as this constitutes the barrier for declaring a molecule RNaseR-resistant. The overlapping area of the linear and circular distributions may originate from RNaseR-resistant linear RNA or from real circRNAs with some sensitivity to the RNaseR treatment.

### SRCP identifies the largest repertoire of *bona fide* circRNAs without false positives

Previous work suggested that circRNA junctions found by more pipelines are more likely to be generated from *bona fide* circRNAs [21, 33]. Therefore, we decided to use that criterion for selection of the most appropriate cut-off for the RNaseR/Total ratio. Briefly, we looked at the proportion of validated and false-positive circRNAs when circRNAs were identified by one to five pipelines. As we wanted to choose a reasonable criterion for distinguishing *bona fide* from false-positive circRNAs, we looked at this parameter as we changed the cut-off value (Table 1). We observed that by using a cut-off of 0.95 (5% error, a cut-off that includes 5% of linear mRNAs), we included most circRNAs identified by all pipelines (while eliminating 95% of linear RNAs). Below 5% error, the proportion of the true circRNAs selected decreased rapidly even for those circRNAs detected by all the pipelines, which are highly likely to be real. On the other hand, higher error rates result in the inclusion of false positives (i.e., as the cut-off was made less stringent; Figure 2C). This was also the trend when we examined circRNAs detected by individual pipelines (Figure S1). Using this criterion (5% of false positives) we included most real circRNAs, while minimizing the number of false positive.

**Table 1:** Cut-off and performance.

| Number of pipelines<br>that identified the<br>putative circRNA | Cut-off |  |  |  |  |  |
| --- | --- | --- | --- | --- | --- | --- |
|  | 0.9 |  | 0.95 |  | 0.99 |  |
|  | TRUE | FALSE | TRUE | FALSE | TRUE | FALSE |
| 1 | 35% | 65% | 27% | 73% | 8% | 92% |
| 2 | 51% | 49% | 41% | 59% | 19% | 81% |
| 3 | 80% | 20% | 70% | 30% | 39% | 61% |
| 4 | 88% | 12% | 77% | 23% | 56% | 44% |
| 5 | 97% | 3% | 92% | 8% | 77% | 23% |

By utilizing the SCRP strategy with a cut-off of 5%, the proportion of circRNAs identified increased from about 10% (~1900 circRNAs; Figure 2A) to about 56% (~2800 circRNAs; Figure 2D). Importantly, some pipeline-specific circRNAs were identified as true circRNAs by the SCRP approach. Most of these pipeline-specific circRNAs have an RNaseR to total RNA ratio lower than the cut-off value (Figure 2E).

Using the chosen threshold, we determined how well each pipeline annotated circRNAs. We identified false positives and false-negative circRNAs based on RNaseR susceptibility. The proportion of true positives was nearly the same for all pipelines (~50%); however, the percentages of false positives and false negatives varied across pipelines and, in most cases, exceeded the number of identified *bona fide* circRNAs (Figure 2F). These results demonstrate that the utilization of multiple pipelines and analysis of RNaseR susceptibility as implemented in SCRP is more accurate than any previously described pipeline for annotating circRNAs.

### Expression and genomic features of *bona-fide* circRNAs

Given our identification of a set of *bona fide* circRNAs, we asked whether these circRNAs could be identified using other genomic and/or expression features. As circRNAs lack a polyA tail, their appearance in polyA^+^ libraries is usually an indicator of a false positive (the exceptions are the few circRNAs that contain stretches of adenosine within them). Indeed, validated circRNAs had very low expression levels in polyA^+^-selected RNA-seq libraries (Figure 3A). In addition, *bona fide* circRNAs tend to be expressed at higher levels and be longer than false-positive circRNAs identified in total RNA-seq libraries (Figure 3B, C). These two features could potentially be useful in identifying circRNAs.

**Figure 3.**
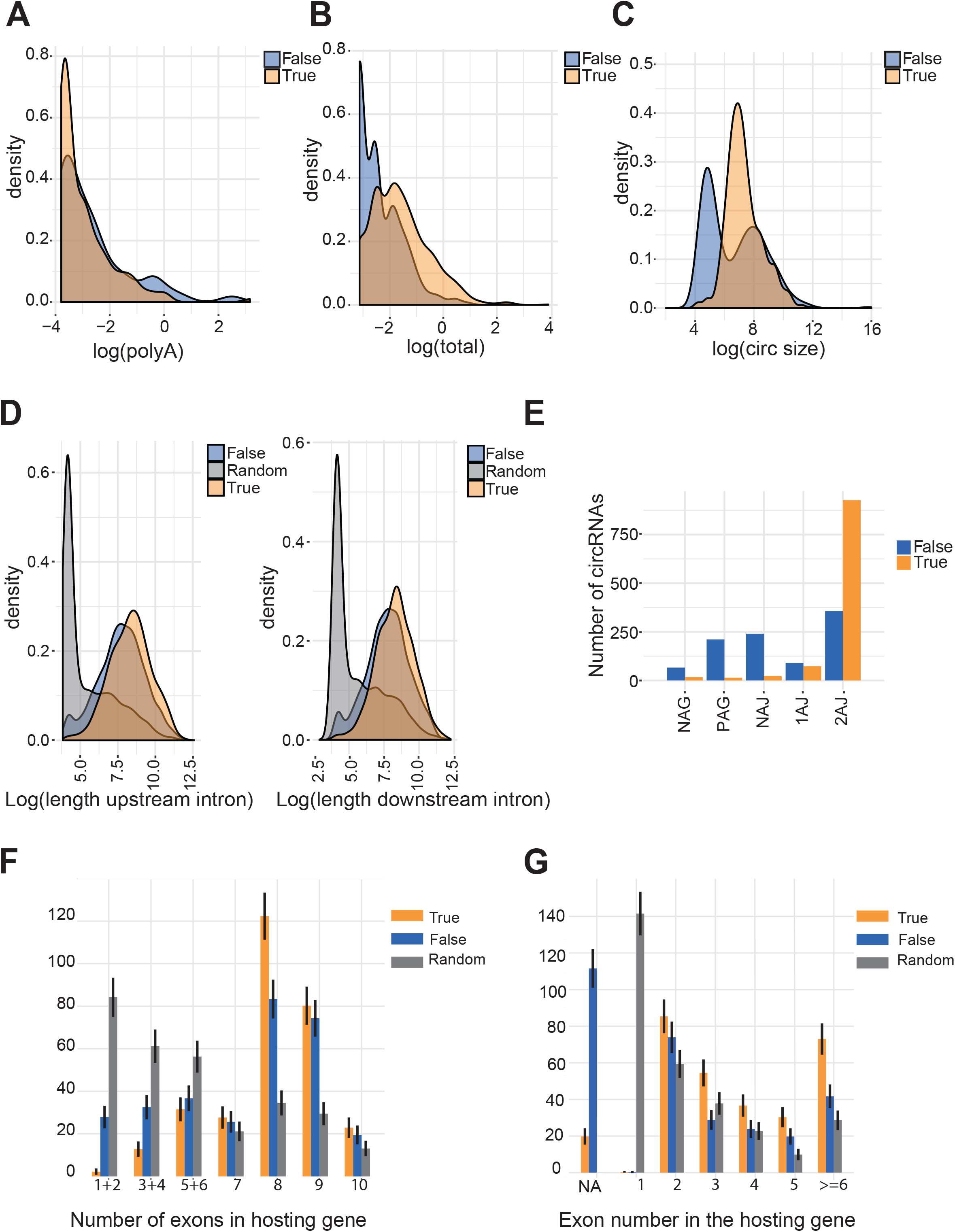
Certain genomic features distinguish circRNAs. **A.** PolyA distribution of the true-positive and false-positive circRNAs in a *D. melanogaster* library. **B.** Expression distribution of true-positive and false-positive circRNAs. **C.** Distribution of the circRNA size in true-positive and false-positive circRNAs. **D.** Distributions of the intron lengths flanking true-positive and false-positive circRNAs and randomly selected exons in the upstream direction **(left)** and downstream direction **(right)**. **E.** Number of indicated annotations for exons of true-positive and false-positive circRNAs and randomly selected exons. **F.** Number of exons in the hosting gene for true-positive and false-positive circRNAs and randomly selected exons. **G.** Numbers of exons true-positive circRNAs, false-positive circRNAs, and randomly selected exons that are the second exon of the hosting gene.

To identify additional genomic feature differences between circRNAs and linear transcripts, we compared circRNAs to a group containing exons randomly selected from the group of exons that do not form circRNAs (see Methods). Neither intron length nor GC content clearly discriminated between true and false-positive circRNA junctions, although exons included within true and false-positive circRNA junctions were flanked by much longer introns than randomly selected exons (Figure 3D) in agreement with an analysis performed previously [38]. In addition, the exons on both sides of *bona fide* circRNAs tended to be annotated (Figure 3E). Hence, many of the junctions that are wrongly classified as circRNA-specific are generated from poorly annotated genes. Interestingly*, bona fide* circRNAs tended to be hosted by genes with more exons than false-positive circRNAs or randomly selected exons (Figure 3F). As previously described [39], *bona fide* circRNAs were more likely to be generated from the second exon of the hosting gene than were randomly selected exons, although we observed a similar trend for the false-positive candidates (Figure 3G). Thus, *bona fide* circRNAs have genomic features that distinguish them from exons that are not circularized, but these differences are not enough to design a non-experimental criterion for the identification of “true” circRNAs from RNA-seq data.

### Annotated circRNAs can be accurately quantified using SRCP

circRNA pipelines identify different sets of circular junctions. As these pipelines utilize different quantification approaches, their results cannot be merged and compared for downstream analysis. The methodology described here overcomes this limitation. SRCP merges potential circRNA junctions identified using various pipelines and then quantifies them using a seed-matching approach. This allows calculation of differential expression of all *bona fide* circRNAs, independently of which circRNA pipelines detected them. Theoretically, the SRCP performance should be strongly affected by the length of the seed utilized to identify the circRNA junctions. Longer seeds should unequivocally identify the circRNA junction, and shorter seeds should result in higher false-positive rates. We ran the SRCP with three different seed lengths; 6, 8, and 10 bases. As expected for each library depth, as we shortened the seed we detected more circular reads and a larger repertoire of circRNAs (Figure S2A). Changing the seed length while keeping the read length and library depth constant only marginally altered the circRNA detection, although seed length was more important for shorter RNA-seq reads than longer (Figure S2B).

To determine how the quantification varies with the depth of the RNA-seq dataset and lengths of reads, we utilized a previously published fly head dataset (SRR1197359). We first randomly down-sampled the data to obtain subsamples with 10 million, 30 million, and 50 million reads and determined the accuracy and sensitivity of the different pipelines and our SRCP approach. As expected, with higher depth more actual circRNAs were detected by all pipelines (Figure 4A). We obtained similar results when we focused only on the 100 circRNAs with high levels of expression (Figure S3A). These results demonstrate the accuracy of SRCP for circRNA quantification.

**Figure 4.**
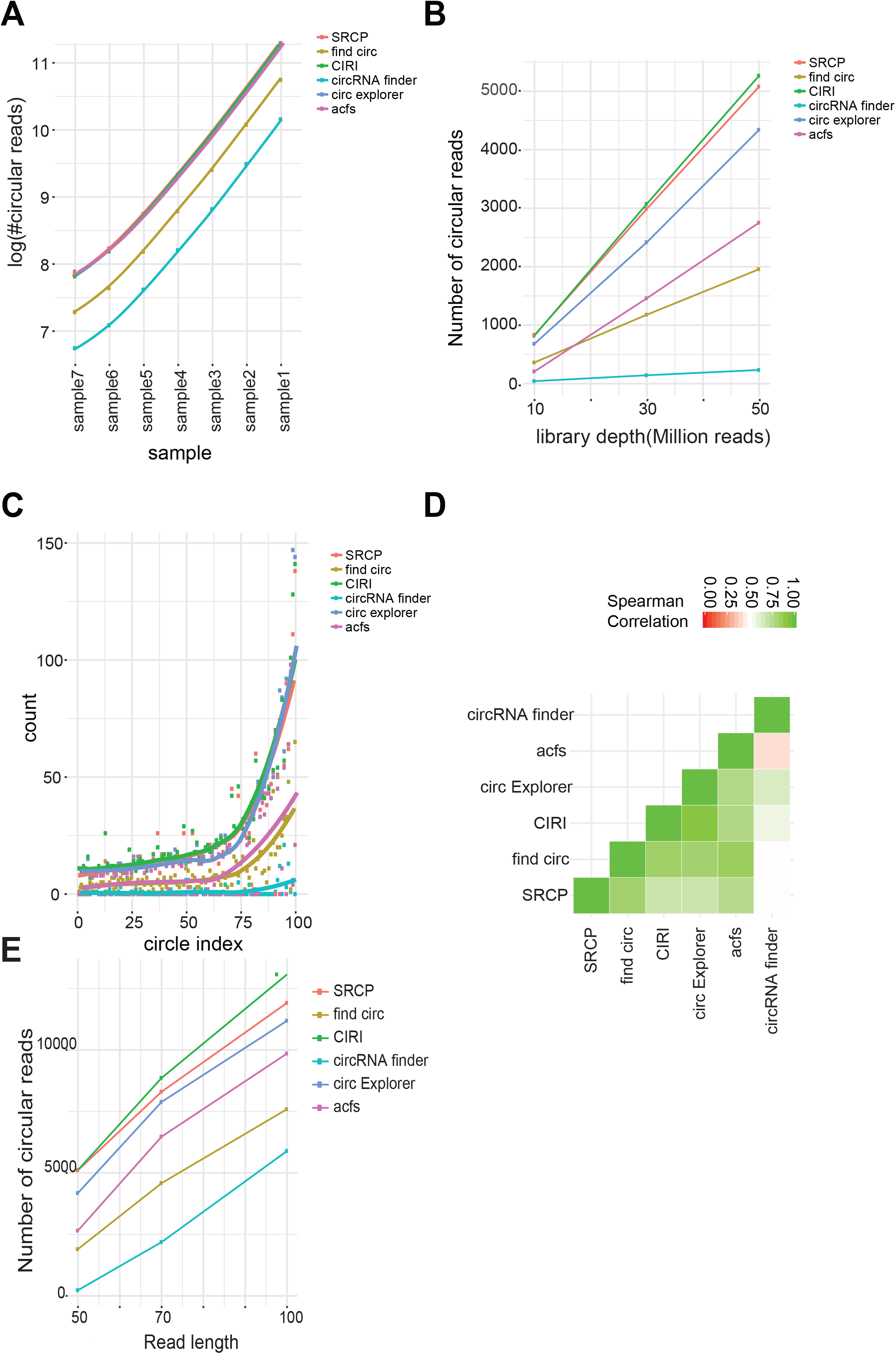
circRNAs can be accurately quantified using seed matching. **A.** Number of circRNA reads for true-positive circRNAs detected by all pipelines **B.** Number of circRNA-associated reads found by each circRNA-identification pipeline and by SRCP as a function of read depth **C.** Number of circRNA reads found for the 100 circRNAs expressed at the highest levels. **D.** Spearman correlation heatmap visualizes quantification by the different pipelines and SRCP. **E.** The number of circRNA reads found in each sample by SRCP using a seed of length 4.

**Figure 5.**
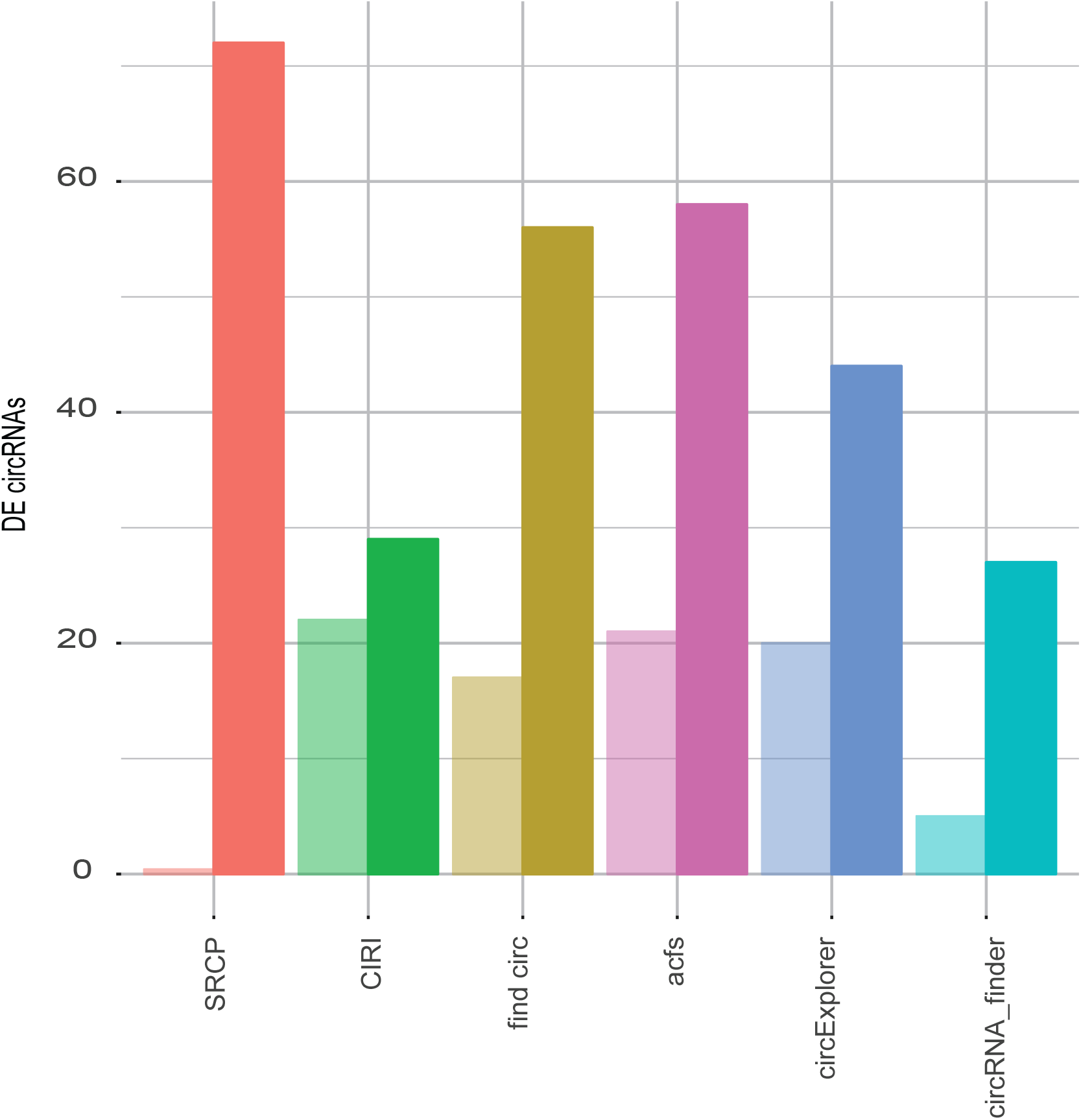
SRCP enables accurate identification of more differentially expressed circRNAs than other pipelines. Number of true (darker bars) and false-positive (lighter bars) differentially expressed circRNA found by SRCP and the circRNA-identification pipelines.

To determine the accuracy and sensitivity of our seed-based quantification we used both simulated and experimental data. First, we simulated seven sets of RNA-seq reads based on known circRNA junctions from *Drosophila melanogaster* (see Methods). Each set was designed to contain half of the number of circRNA read counts as the one before (Table 2). Importantly, these simulated data utilized circRNA junctions that were identified by all pipelines, and hence the results reflect only the quantification aspects of the pipelines. Of the pipelines tested SRCP, CIRI, circExplorer, and acfs detected the most circRNA reads in each sample with very small variations; find_circ and circRNA_finder were less sensitive (Figure 4B). This means that SRCP, CIRI, circExplorer, and acfs are sensitive enough to detect low abundance circRNAs represented by only 10 to 40 reads per circle per sample (Table 2, samples 5-7).

**Table 2:** Simulated samples and number of reads.

| Sample | batch1 | batch2 | batch3 | batch4 | batch5 | batch6 –<br>control |
| --- | --- | --- | --- | --- | --- | --- |
| 1 | 1000 | 800 | 600 | 400 | 200 | 50 |
| 2 | 500 | 400 | 300 | 200 | 100 | 50 |
| 3 | 250 | 200 | 150 | 100 | 50 | 50 |
| 4 | 125 | 100 | 75 | 50 | 25 | 50 |
| 5 | 63 | 50 | 37 | 25 | 13 | 50 |
| 6 | 32 | 25 | 19 | 13 | 7 | 50 |
| 7 | 16 | 13 | 10 | 7 | 4 | 50 |

SRCP, circExplorer, and CIRI identified more *bona fide* circRNAs than acfs, find_circ, and circRNA_finder at all depths tested (Figure 4A). This is because the later three pipelines identify many reads as circRNAs that are actually false positives (Figure 2F). When sensitivity was evaluated, SRCP, CIRI and circExplorer again out-performed the other methods (Figure 4B). When we evaluated the number of circRNA reads detected for the 100 most highly expressed circRNAs, we observed similar performances for SRCP, CIRI, circExplorer, and acfs. These pipelines identified significantly more junctions than find_circ and circRNA_finder (Figure 4C). These results demonstrate that the use of SRCP guaranties no false positives with sensitivity similar to that of the more sensitive circRNA-detection pipelines.

We next quantified the circRNAs found in the fly head data [40] using the different pipelines and SRCP and compared the results. The SRCP quantification best correlated with quantifications from find_circ and acfs pipelines and least well with circRNA_finder (Figure 4D). As most pipelines rely on the identification of hybrid RNA-seq reads that align to two regions of the transcriptome, circRNA identification and quantification becomes less efficient as RNA-seq reads become shorter. In theory, SRCP should not have this problem as once indexes are established the quantification of circRNAs relies on 10 bases or less. To test this, we used an evaluated subsample of 50 million reads with different lengths (50 bases, 70 bases and 100 bases). We found that SRCP outperformed the other pipelines except for CIRI (Figure 4E). We also see that SRCP outperforms the other pipelines while the subsamples are of different depth Figure 4B. These results suggest that SRCP could be also used to quantify circRNAs from short RNA-seq libraries like those generated from RNA precipitation-based methods such as RIP and CLIP and other techniques. Only SRCP and CIRI will be effective with techniques that generate very short RNA-seq reads like ribosome footprinting for which we utilized a similar approach [16].

### SRCP identifies more differentially expressed circRNAs than individual pipelines

Using DESeq2 [41], we looked for differentially expressed circRNAs. We found that the SRCP pipeline more effectively detected differentially expressed circRNAs than all other pipelines (Figure S4). Importantly, the simulated data contained circRNAs that were detected by all the pipelines, which favors individual pipelines (in particular CIRI2, which had a higher number of false positives but the best quantification power). Therefore, we performed the same analysis with previously analyzed *Drosophila* datasets (SRR1197279, SRR1197275, SRR1197273, SRR1197274, SRR1197472, SRR1197473, SRR1197474, SRR1197362). In these datasets, we sought to identify the circRNAs differentially expressed in young (1 day old) versus aged (20 days old) flies. Using the SRCP approach, we identified 63 *bona fide* circRNAs that are differentially expressed (FDR<0.05) between these two conditions; other pipelines identified fewer (Figure 6).

Importantly, using each of the pipelines without pre-filtering for true-positive junctions resulted in sets that contained more differentially expressed circRNAs but that included many false positives (Figure 6, Table 3). For example, find_circ identified 74 differentially expressed circRNAs, 57 (77%) of them involve *bona fide* circRNAs and 17 (23%) false positives. These results emphasize the importance of the annotation step that we present here. Using our annotation procedure lowers the number of false positives and thus results in higher accuracy in further analyses. In sum, these results show that the use of our two-step procedure extracts more accurate information from the RNA-seq data than other circRNA identification pipelines.

**Table 3:**
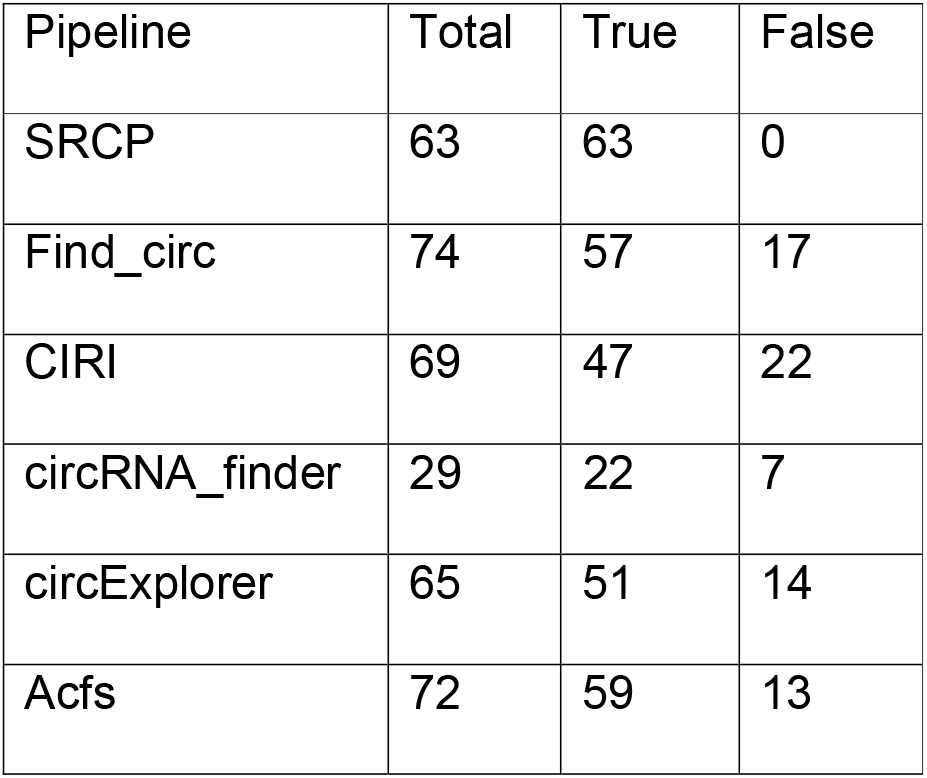
Differentially expressed circRNAs identified using SRCP and different circRNA-identification pipelines.

| Pipeline | Total | True | False |
| --- | --- | --- | --- |
| SRCP | 63 | 63 | 0 |
| Find_circ | 74 | 57 | 17 |
| CIRI | 69 | 47 | 22 |
| circRNA_finder | 29 | 22 | 7 |
| circExplorer | 65 | 51 | 14 |
| Acfs | 72 | 59 | 13 |

## DISCUSSION

In this study, we present a novel computational pipeline that facilitates accurate annotation and quantification of circRNAs from RNA-seq data. The method, which we call SRCP, quantifies circRNAs with high sensitivity and without false negatives. The method is general and is not limited to specific circRNA detection pipelines. The quantification step it is faster and simpler than all other pipelines since it does not search *de novo* for circular RNA junctions.

A number of circRNA identification and quantification tools have been described including Acfs [24], DCC [25], segemehl [26], CIRCexplorer [27], KNIFE [28], MapSplice2 [29], circRNA_finder[30], CIRI [31], and find_circ [11]. All can be used to identify and quantify circRNAs, but none of identify all circRNAs. For example, only about 10% of circRNAs from the dataset analyzed here were identified by all pipelines. Therefore, when analyzing samples for differentially expressed circRNAs researchers generally chose one pipeline or focus on those circRNAS detected by multiple pipelines. This is unfortunate as this procedure eliminates many real circRNAs from the analysis. Moreover as different pipelines utilize different quantification approaches their results cannot be combined. SRCP solves this issue by first annotating all circRNAs and then utilizing common criteria for quantification. Our approach thus analyzes a much larger group of circRNAs than would be interrogated using only one method or the overlapping circRNAs of several methods and accurately quantifies circRNAs without false positive or negatives. The approach utilized by SRCP is somehow similar to the one used by KNIFE [28]. However, KNIFE examines and counts all possible splicing junctions, making it slower and very demanding from the computational point of view.

One of the strengths of our approach is that the user can modify the annotation of the false positive circRNAs identified by different circRNA detection pipelines. As the annotation is based on the RNaseR-treated library to the total (rRNA depleted) library ratio, the user can control the false positive rate.

Interestingly, in our analysis of *Drosophila* circRNAs identified and quantified using SRCP, we found that *bona-fide* circRNAs share some genomic features and expression characteristics. These include their very low abundance in polyA-selected RNA-seq libraries and slightly lower expression levels, larger size, hosting by genes with more introns, and better genomic annotation compared to randomly selected exons. Although none of these features individually allowed us to computationally discriminate between “true” and “false” circRNAs, machine learning approaches might be trained to take these features into account to improve *de novo* identification of circRNAs in order to diminish false negatives.

### Conclusions

In sum, SRCP is a novel method that combines circRNA annotation and an efficient algorithm for quantification. To identify false positives from the several different circRNA-identification pipelines, we compared the expression of the putative circRNAs in mock and RNaseR-treated samples. By comparing the results obtained using SRCP with the results obtained using five circRNA identification and quantification pipelines on multiple simulated and real RNA-seq datasets, we found that SRCP identifies more differentially expressed circRNAs than any of the other methods. Furthermore, SRCP allows quantification of circRNAs that were identified by different pipelines.

## METHODS

### Creation of a circular reference

Using available genome annotation, we extract for every circRNA all potential transcripts. Next, we score the transcripts as follows: (i) If the start coordinate and the end coordinate of the circle are both exactly on a 5’ and 3’ boundaries of the transcript’s exons, the score is maximal. (ii) If only one coordinate is exactly on an exon boundary and the other is not, the score is 1. (iii) If neither coordinate is on any exon boundary the score is 0. Next, we choose for each circRNA in the database, a transcript that best fits the circle. This is the transcript with the highest score or, if a few transcripts all have the same highest score, one is randomly chosen. Next, using bedtools getfasta [42] we extract the circle sequences of the chosen transcript or transcripts and create an index using bowtie2 [43]. This is done for each potential circRNA.

### Creation of a linear index

In order to create linear references for each circle junction, we look for the closest annotated exons upstream and downstream. We then extract the sequences for these exons with bedtools get_fasta [42] and build the index with bowtie2 [43].

### Detection and quantification of circRNAs

RNA-seq reads are aligned to the circular index with bowtie2 [43]. The reads that align to the circular index are next aligned to the genome and to the transcriptome. The alignment to the genome is done in order to remove reads that come from unannotated genes. The alignment to the transcriptome is done to ensure that no reads that are originally from a linear transcript are included in the detection and quantification of circRNAs analysis. Reads that align to the circular index and not to the genome or transcriptome are candidates for circular circRNA reads. To be confirmed as a circRNA, the junction has to be included in the read and a certain number (*j*) bases upstream or downstream or the putative junction have to match the exon with no mismatches.

To quantify circular reads, the number of reads that align to the circular index for each circRNA is calculated. For linear reads, the number of reads that align to the linear index is calculated for all transcripts once for the downstream of the circle and once to the upstream side of the circle.

### Selection of the cut-off value

To select an appropriate cut-off value, we compared the number of circular and linear RNA reads classified as RNaseR-resistant for an array of different cut-off values of the RNaseR/Total ratio (Figure S2A, B). We performed this analysis for all the pipelines utilized in this study including SRCP. We first rescaled the data. The cut-off value is expressed as the percentage of linear RNAs would be considered RNaseR-sensitive (i.e., a 0.90 cut-off value excludes 90% of the linear mRNAs). A cut-off of 95% removes most linear RNA reads (Figure 2E). Interestingly, circRNAs identified by more than one pipeline tended to be enriched among those considered real at higher cut-off values (Figure 2E). For example, a cut-off of 0.8, 98% of the circRNAs that were found by all pipelines (and are believed to be true circRNAs) were annotated as *bona fide* circRNAs. On the other hand, at a cut-off of 0.99 only 77% were found by all five pipelines to be *bona fide* circRNAs. At a cut-off of 0.95, 92% were *bona fide* circRNAs (Figure 2E, Table 1). Thus, the lower the cut-off, the more false positives we introduce. Elevating the cut-off results in fewer false positives but also fewer true positives.

### Creation of simulated data

In order to create a simulated dataset we took all the circular junctions found in *Drosophila melanogaster* heads and extracted the fasta sequences for the entire circle plus the fasta sequences for the upstream and downstream exons. We then randomly selected start positions on the circRNA and generated 70-base circular reads (from the circle sequences) and linear reads from the upstream and downstream exons.

### Selection of random exons

To select random exons, we used the *Drosophila melanogaster* annotation from UCSC. For each gene we selected the starting exon in the transcript. We then randomly selected the desired number of exons and created a new annotation for each of these transcripts.

### Detection and quantification of circRNAs with the different pipelines

We ran circRNA detection pipelines Acfs, CircExplorer, circRNA_finder, CIRI and find_circ with default parameters with the dm3 genome and genome annotation from the UCSC genome browser as input.

### Differential expression analysis

We normalized the circular reads to the number of aligned reads in the sample. We used DESeq2 [41] to detect the differentially expressed circs. We combined the results from all pipelines into one data frame and filtered out circRNAs that had less than two reads. We then performed the analysis and selected the circRNAs with adjusted p value < 0.05 as significantly differently expressed.

## List of abbreviations

circRNA: circular RNAs
CNS: central nervous system
SRCP: Short Read CircRNA Pipeline
GEO: Gene Expression Omnibus
SRA: Sequence Read Archive

## Declarations

### Availability of data and material

SRCP is implemented in python. Its source code is available at: https://github.com/avigayel/SRCP/. The datasets analyzed are available in the GEO and SRA databases.

### *Drosophila melanogaster* datasets

total and RNaseR fly heads at 18 °C and 29 °C: GSM1347830, GSM1347831, GSM1347838, GSM1347839, GSM1347842, GSM1347843, GSM1347834, GSM1347835.

### Fly aging dataset (for quantification analysis)

fly heads at age 22 days: SRR1197359.

### Fly aging dataset for differential expression analysis

4 samples of 1-day-old female fly heads plus 4 samples of 22-day-old female fly heads: SRR1197279, SRR1197275, SRR1197273, SRR1197274, SRR1197472, SRR1197473, SRR1197474, SRR1197362.

### Simulated dataset

created *in silico* using the junctions that were detected from fly heads in total and RNaseR libraries: GSM1347830, GSM1347831, GSM1347838, GSM1347839, GSM1347842, GSM1347843, GSM1347834, GSM1347835.

## Ethics approval and consent to participate and consent for publication

Not applicable.

## Author contributions

S.K. conceived and designed the project. A.R. collected and analyzed the data with the help from D.A. and R.A.-F. S.S. built the structure of the pipeline. S.K. and A.R. wrote the paper with input from all authors.

## Acknowledgements

We thank members of the Kadener lab for comments.

## Supplementary figure legends

**Figure S1.**
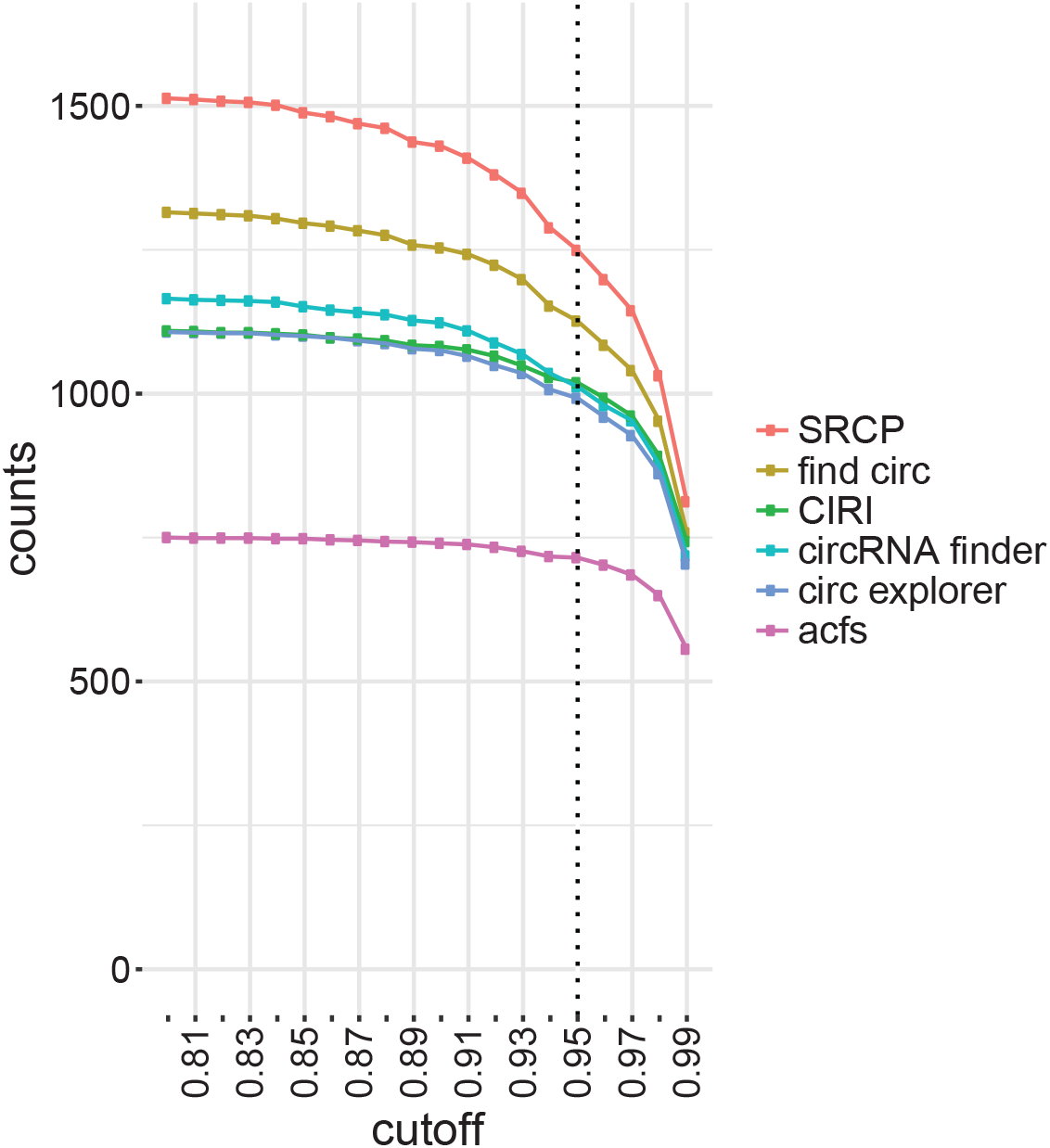
Removing the false positives circRNAs. Number of true-positive circRNAs identified by each pipeline using different cut-off values.

**Figure S2.**
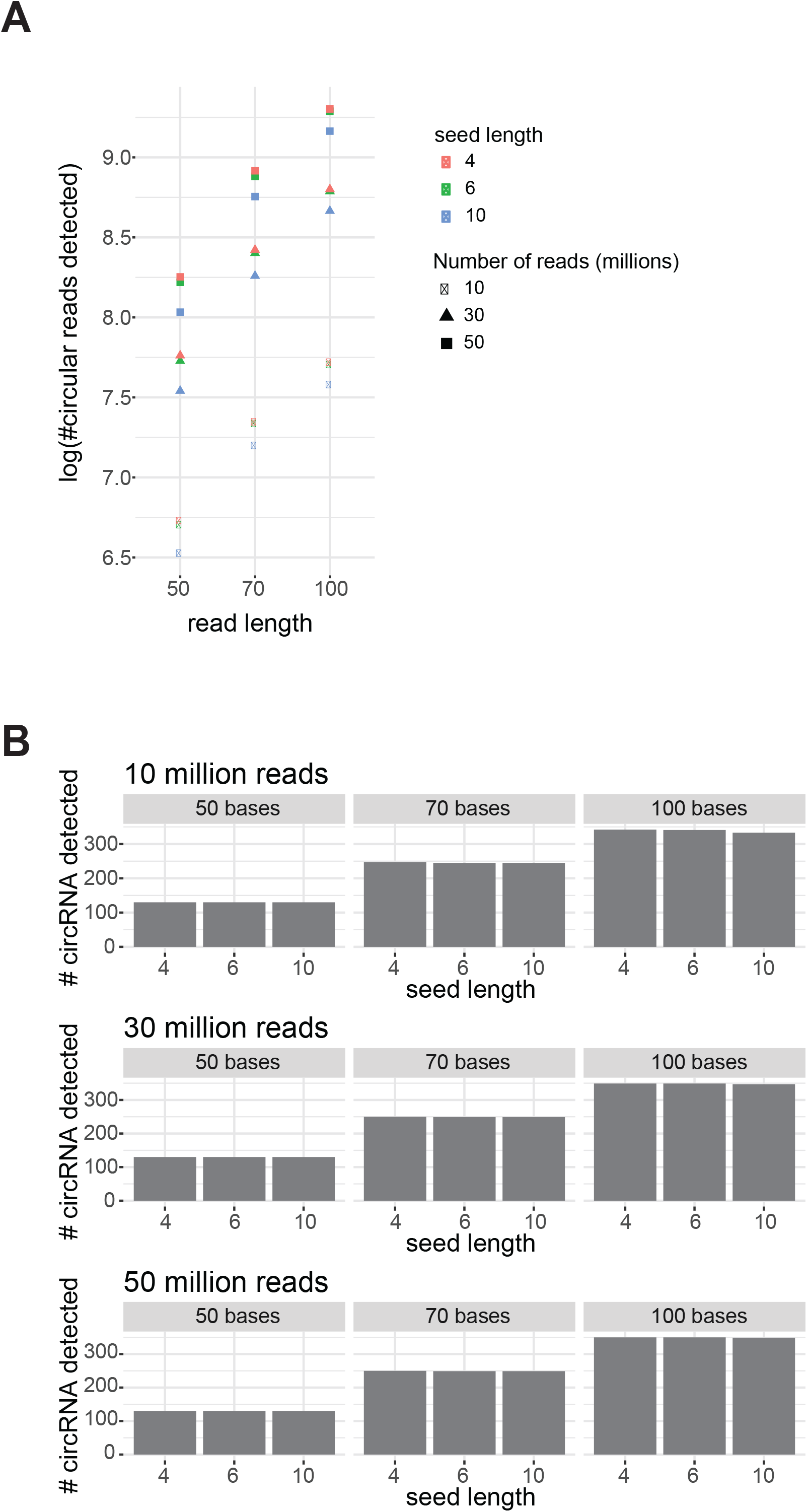
SRCP detects circular reads in samples with low coverage depth. **A.** Number of circular reads detected by SRCP for the different read lengths and sample depths. **B.** Number of circRNAs detected in the different samples using different lengths of seeds.

**Figure S3.**
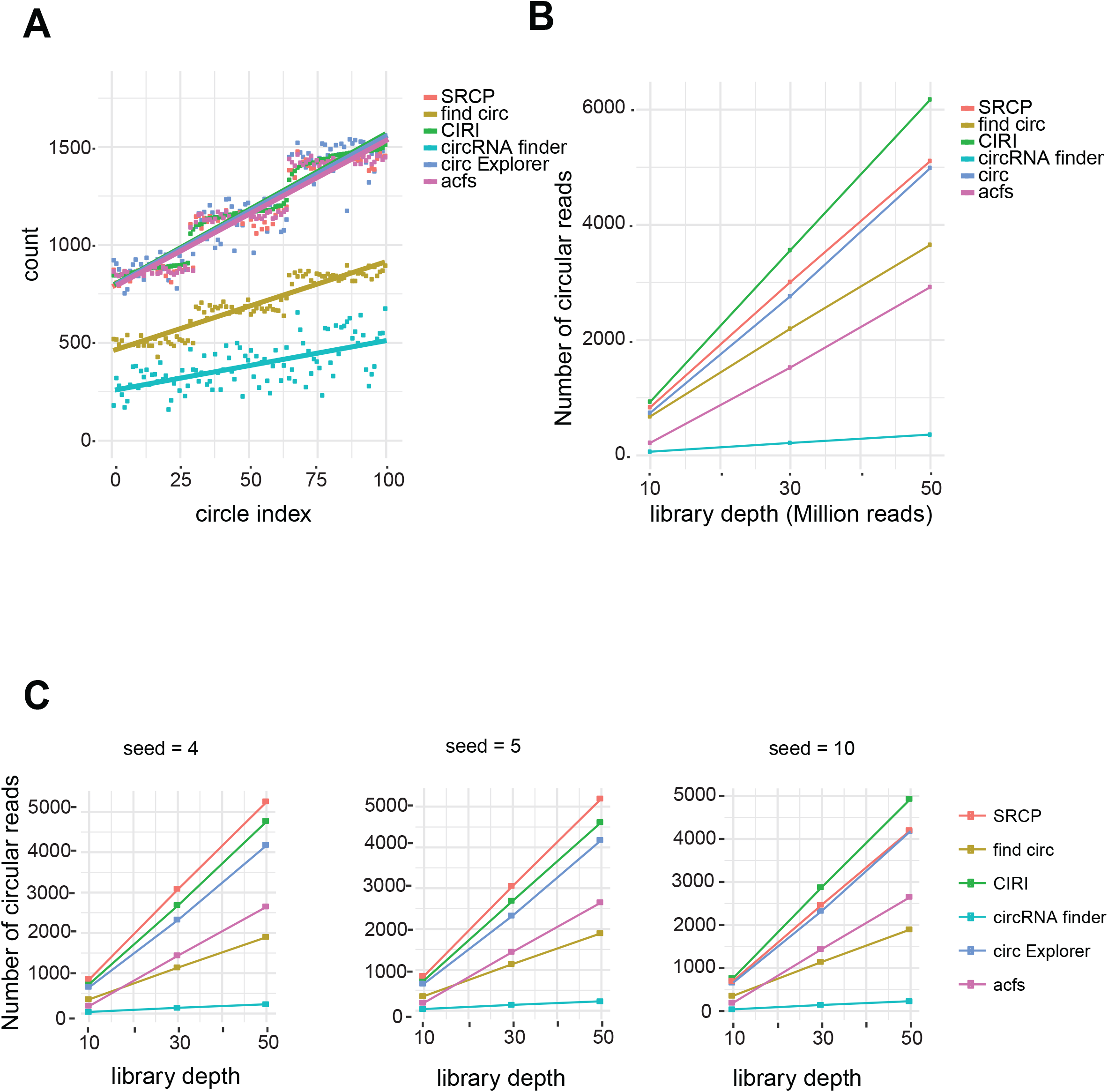
Simulated circRNAs can be accurately quantified using seed matching. **A.** Number of circRNA reads found by SRCP and circRNA-identification pipelines in each sample. **B.** Number of circRNA reads found by each pipeline for the 100 most highly expressed circRNAs in the simulation. **C.** Number of circRNA reads found in the different samples using different seeds.

**Figure S4.**
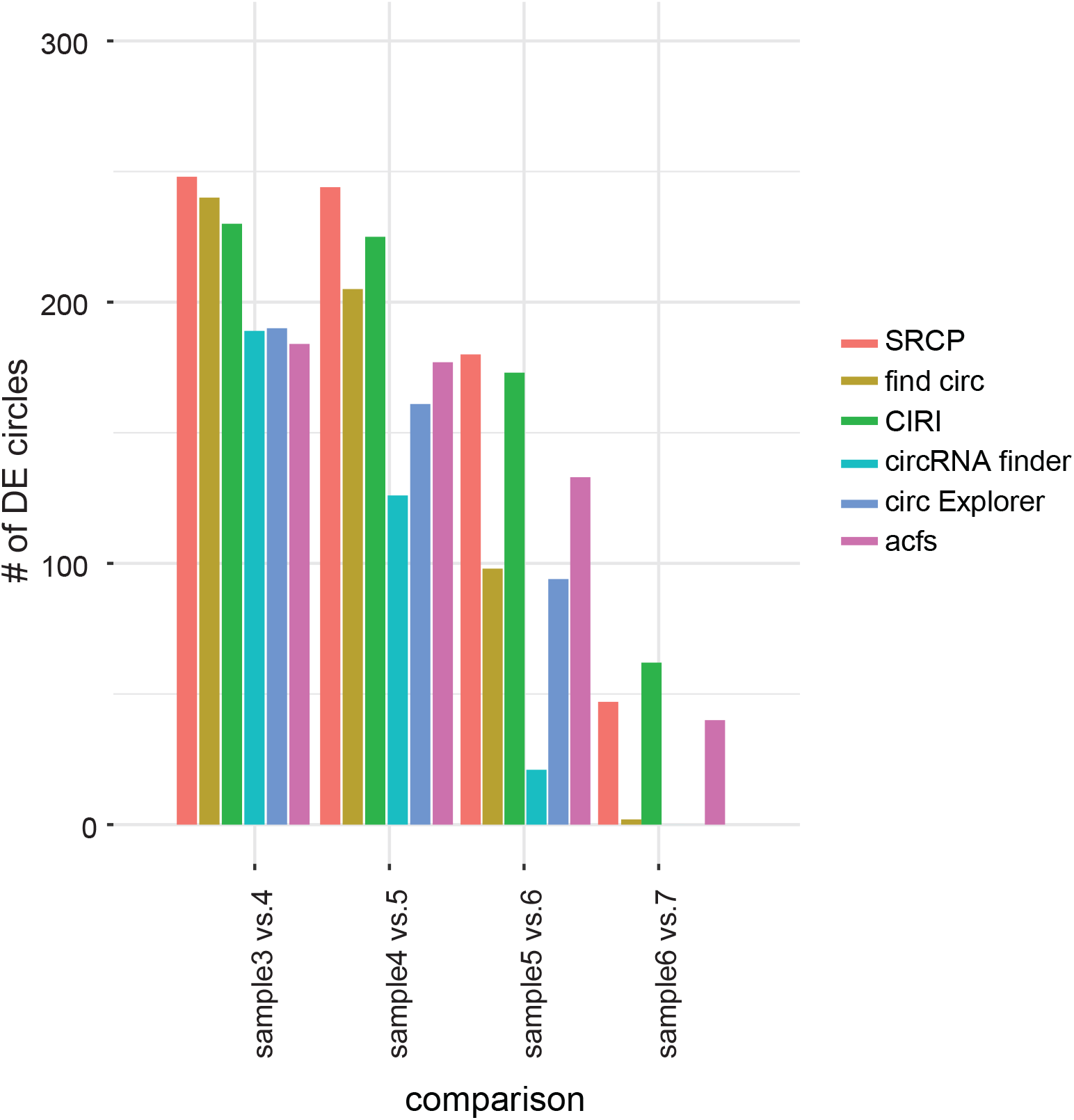
SRCP detects more simulated differentially expressed circRNAs than circRNA-identification pipelines. Number of differentially expressed reads found by SRCP and each pipeline in the different sample comparisons.

